# Expectations about presence enhance the influence of content-specific expectations on low-level orientation judgements

**DOI:** 10.1101/2024.02.22.581334

**Authors:** Joost Haarsma, Aaron Kaltenmaier, Stephen M Fleming, Peter Kok

## Abstract

Will something appear and if so, what will it be? Perceptual expectations can concern both the presence and content of a stimulus. However, it is unclear how these different types of expectations interact with each other in biasing perception. Here, we tested how expectations about stimulus presence and content differently affect perceptual inference. Across separate online discovery (N=110) and replication samples (N=218), participants were asked to judge both the presence and content (orientation) of noisy grating stimuli. Crucially, preceding compound cues simultaneously and orthogonally predicted both whether a grating was likely to appear as well as what its orientation would be. We found that expectations of presence interacted with expectations of content, such that the latter’s effect on discrimination was larger when a stimulus was expected to appear than when it was not. This interaction was observed both when a grating was truly presented and when participants falsely perceived one. Confidence in having seen a grating on the other hand was independently affected by presence and content expectations. Further, modelling revealed higher sensitivity in distinguishing between grating presence and absence following absence cues than presence cues, demonstrating an asymmetry between gathering evidence in favour of stimulus presence and absence. Finally, evidence for overweighted predictions being associated with hallucination-like perception was inconclusive. In sum, our results provide nuance to popular predictive processing accounts of perception by showing that expectations of presence and content have distinct but interacting roles in shaping conscious perception.

## 1 Introduction

The sound of an opening door elicits the expectation of something entering through its frame. Turning our gaze towards the door, sensory evidence then either affirms or denies this expectation. The predictive processing (PP) framework postulates that this process is inherent to all of perception such that what we perceive represents a combination of predictions, elicited by cues for example, and incoming sensory information (Bastos et al., 2012; Friston, 2005; Hohwy & Seth, 2020). Indeed, empirical research in recent decades suggests that predictions can bias perception towards the expected outcome through top-down connections that descend along the neural processing hierarchy (Aitken, Turner & Kok, 2020; Brandman & Peelen, 2017; Chalk et al., 2010; de Lange, Heilbron & Kok, 2018; Haarsma et al., 2022; Haarsma et al., 2023; Kok et al., 2013; Summerfield & de Lange, 2014; Thomas et al., 2022; Wyart, Nobre & Summerfield, 2012; Yon et al., 2018).

Importantly, however, not all predictions are the same. Consider the earlier example of expecting something or someone to enter the room when hearing a door open. Rather than expecting a specific content, such as a specific person or cat, the sound of the door elicited a prediction of general stimulus presence – something being there. However, based on the time and place, expectations are likely also held regarding the nature of the stimulus entering the room, after all a colleague is far more likely to enter your office than a cat. How then do these different kinds of perceptual predictions, those about stimulus presence and those about stimulus content, combine to shape perception? This question has further implications to predictive processing accounts of psychiatric disorders characterized by aberrations in perception such as autism (Lawson, Rees & Friston, 2014, Sapey-Triomphe et al., 2023) and psychosis (Corlett et al., 2019, Powers et al., 2017). Specifically in the latter, previous studies have found that perceptual predictions can particularly affect perception in those prone to hallucinations, and even induce *de novo* percepts (Haarsma et al., 2020; Schmack et al., 2021; Sterzer et al., 2018; Teufel et al., 2015). Yet, these studies typically do not dissociate predictions about presence and contents. Indeed, in a previous study of ours, hallucination proneness was not related to content-based cues. Instead, the content of false percepts arose from activity in the feedforward layers of the early visual cortex, independently of the cues signalling the most likely stimulus. However, implicit beliefs about stimulus presence rather than content could still have contributed to these false percepts (Haarsma et al., 2023). This raises the possibility that predictions of stimulus presence and content could have distinct effects on perceptual inference.

The higher-order state space (HOSS) model attempts to account for this distinction by distinguishing between predictions about stimulus presence (i.e. detection) and its possible contents (i.e. discrimination) at different levels of a processing hierarchy (Fleming, 2020). The result is a hierarchical process in which two distinct perceptual inferences on detection and discrimination are scaffolded by two types of predictions: those about the contents of a stimulus and those about the presence of a stimulus as a whole.

In the present study, we studied how predictions about presence and content interact in altering perception. We orthogonally and simultaneously manipulated expectations of stimulus presence (grating vs. noise) and content (grating orientation) using compound cues. Participants performed a discrimination task on grating orientation, while also indicating their confidence in stimulus presence. The objective of the study was twofold. First, we aimed to establish empirically whether the different types of predictions have distinguishable effects on detection and discrimination responses. Second, considering hallucinations as the result of overweighted predictions, we asked whether presence or content cue effects were differentially related to hallucination-susceptibility, which was assessed using the Cardiff Abnormal Perception Scale (CAPS). To preview, presence predictions moderated the effect of content predictions on discrimination decisions. Meanwhile, presence and content predictions independently influenced stimulus detection. Our initial findings of evidence for a relation between hallucination-proneness and the effect of presence priors on perception did not replicate in a second sample.

## 2 Methods

### 2.1 Participants

Two separate samples were collected with identical paradigms and procedures in order to discover and replicate a possible relationship between prediction effects and hallucination-susceptibility. The first sample consisted of 110 participants (mean age = 28.2, SD = 5.1) and the replication sample contained 218 (mean age = 27.1, SD = 6.0). Participants were recruited through the online recruitment platform Prolific and compensated at a rate of £7.50 per hour. The replication sample size was determined by a power calculation based on the discovered effect size of the relationship between the presence cue effect and hallucination-susceptibility. Participants were required to be fluent in English and between 18 and 35 years old. Data were collected through the online experiment builder Gorilla (Gorilla Experiment Builder, 2022), which was also used to implement the behavioural paradigm. Participants were provided with an information sheet and consent form which they agreed to before moving onto the questionnaires and task. The entire study took approximately one hour and was approved by the local UCL Research Ethics Committee (UCL REC) under the minimum risk ethics protocol for online studies 6649/004.

### 2.2 Task Design & Stimuli

A novel task was created that allowed for the simultaneous manipulation of both stimulus presence and content, inspired by Dijkstra and colleagues (2023). The basic paradigm was adopted from the previously mentioned study by Haarsma et al. (2023) and required participants to discriminate the orientation (left-tilted, 45° or right-tilted, 135°) of a grating as well as report their confidence in having seen a grating at all as opposed to just noise. As such, this confidence response related to a grating’s visibility and should not be confused with metacognitive confidence in the aforementioned discrimination decision. Compound cues were implemented along with other adjustments using Gorilla’s Experiment Builder (Gorilla, 2022).

Following a fixation cross, each trial consisted of a visual cue followed by the target stimulus of either a grating or noise (50% each). Sinusoidal grating patches (spatial freq. = 0.5 cpd) were created using the Psychophysics Toolbox as implemented in MATLAB (Brainard, 1997; The Mathworks Inc, 2022) and combined with noise patches (see below for details on the noise), resulting in gratings with four, five or six percent contrast. These contrast levels were chosen as pilot testing had shown that they result in an adequate task difficulty in most participants. The four noise patches (20% contrast) were created by applying a Gaussian smoothing filter to pixel-by-pixel Gaussian noise. Initially, 1000 noise patches were processed through a bank of Gabor filters with varying preferred orientations. Four noise patches with low (2%) signal energy for all orientations were selected to be included in the present experiment. This assured that the noise matched the gratings in terms of spatial frequency, but without carrying orientation-specific information. These noise patches were used as target stimuli on grating-absent trials, and were presented in counterbalanced manner, to ensure that fluctuations in perception could only result from internal signals, rather than from fluctuations in stimulus noise (Haarsma et al., 2023; Wyart, Nobre & Summerfield, 2012, Pajani et al., 2015). Gratings were tilted either to the left (45°) or the right (135°), again with equal probability. The preceding cues were compounds consisting of a coloured circle predicting the orientation of a possible grating and a surrounding square/diamond predicting the presence of a grating as opposed to noise (see Figure 2, right). The visual cues were created using Adobe Express (Adobe Inc., 2022) and consisted of a coloured circle within a square of a certain rotation as previously explained. Circles were coloured either cyan (HEX: #009999) or orange (#E18000) predicting leftwards (45°) and rightwards (135°) orientations respectively. The surrounding square was black (#000000) and either presented with its horizontal edges in parallel with the horizon or rotated by 45° resulting in a diamond-like shape with the former predicting noise and the latter a grating.

**Fig 1.**
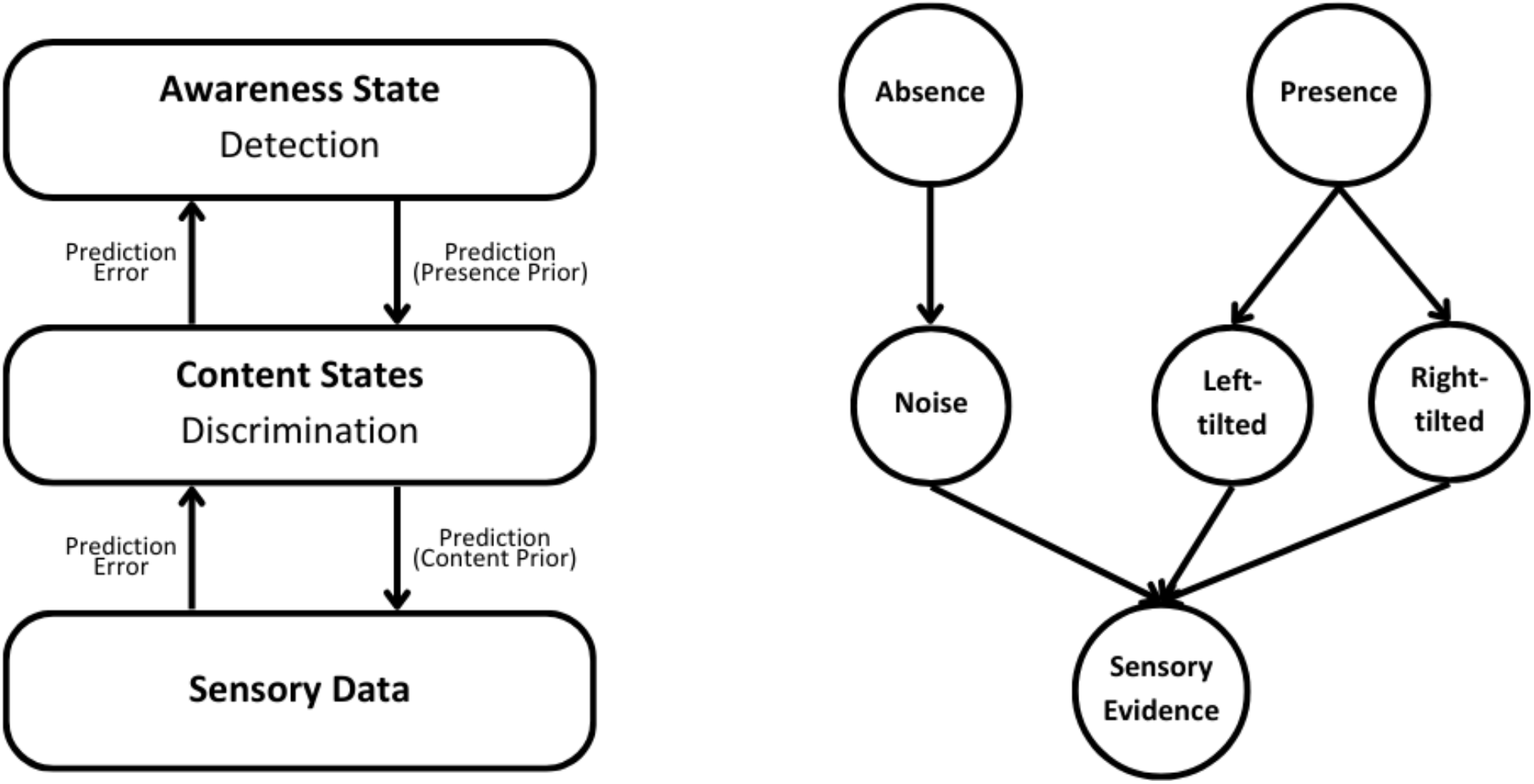
Higher-order State Space Model. Adapted and simplified from Fleming (2020). Tilt in content states refers to gratings in common experimental paradigms.

**Fig 2.**
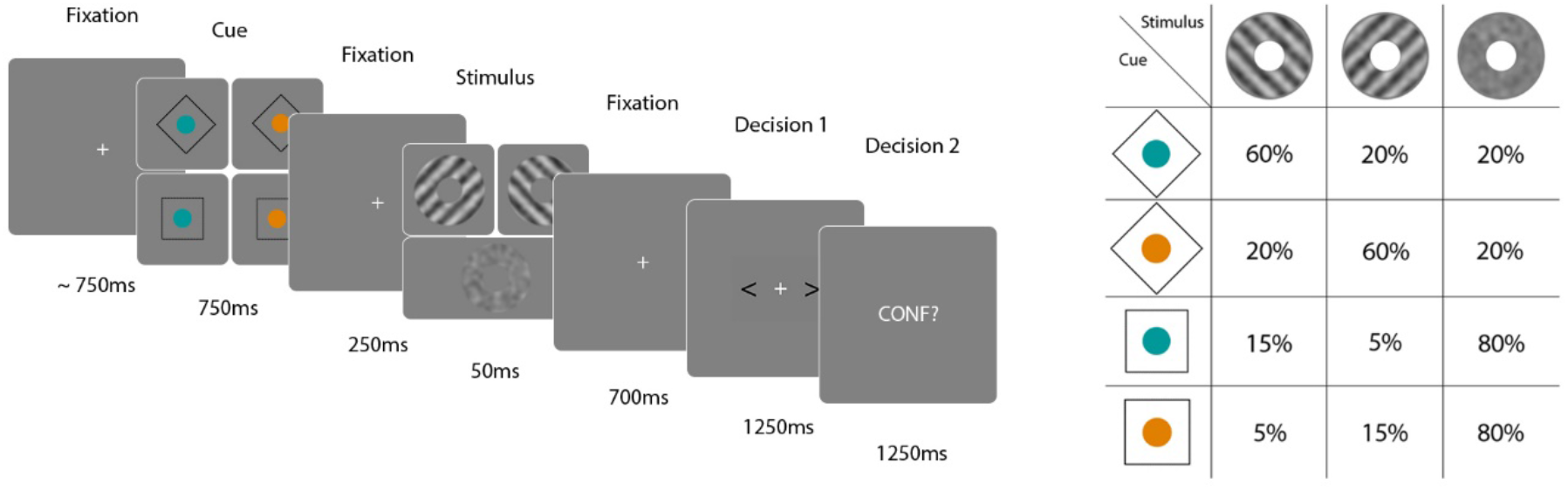
Task design and cue-stimulus-contingencies.

**Figure 3.**
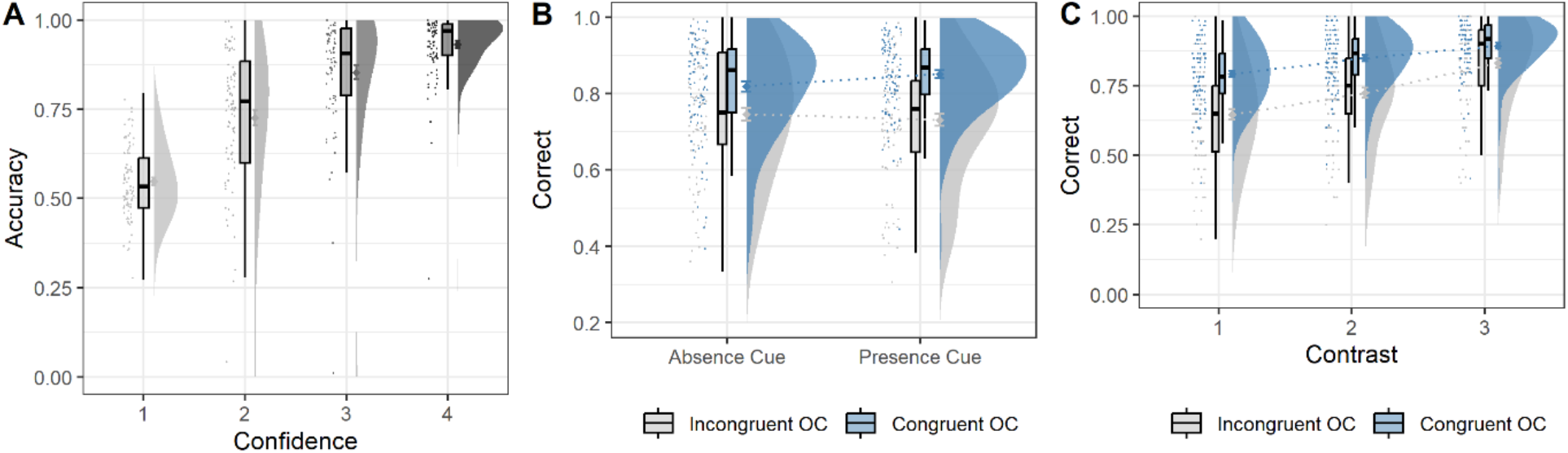
Orientation discrimination accuracy on grating-present trials. (A) Across confidence levels and (B) across cueing conditions (OC = Orientation cue). (C) Effect of orientation cue congruency on accuracy across contrast levels. Error bars reflect within-subject standard error of the mean.

The four different types of cues appeared on an equal number of trials. Crucially, the validity of one cue dimension was independent of the validity of the other cue dimension. In other words, cues indicating a left or right-tilted grating were valid on 75% of trials regardless of whether the cued grating presence was valid. This meant that whether a grating appeared as predicted following a presence cue, or contrary to predicted following an absence cue, did not affect whether the displayed grating would comply with the cued orientation. Conversely, cues indicating grating presence were valid on 80% of trials regardless of whether the cued grating orientation was valid.

Cues were presented for 750ms followed by a 250ms blank screen before the target stimuli appeared for 50ms. Gratings were shown at varying contrast levels as explained in the materials section. Following the target stimulus and another 700ms blank screen, two arrows were displayed, one on each side of a fixation cross. Participants indicated the orientation of a possible grating by selecting the matching arrow on the screen (i.e. the left-pointing arrow for a left-tilted grating response) whose location with respect to the fixation cross varied on a pseudo-random trial-by-trial basis to avoid response biases. Keyboard presses of the buttons ‘U’ and ‘I’ corresponded to picking the arrow on the left and right of the fixation cross respectively. A time-out was recorded after 1250ms. At the end of each trial, participants were shown “CONF?” which prompted them to indicate their confidence in having seen a grating at all. They were asked to do so within 1250ms by pressing one of four keys on their keyboard: Q = “I did not see a grating”, W = “I may have seen a grating”, E = “I probably saw a grating”, R = “I am sure I saw a grating”. The next trial started following a jittered inter-trial interval of either 500, 750 or 1000ms.

### 2.4 Questionnaires

To assess hallucination-susceptibility and control for possible confounds, participants were asked to complete three questionnaires in addition to the described experimental task. Firstly, the Cardiff Abnormal Perception Scale (Bell et al., 2005) was used to measure hallucination-susceptibility consistent with several previous studies in this field. Scores on its three subscales were summed for all 32 items to produce the overall CAPS score for which higher scores indicate higher hallucination-susceptibility. The Beck Depression Inventory was used to assess depression to control for general psychopathology (BDI, Beck, Steer & Brown, 1996). Lastly, a demographics questionnaire was conceived and employed that measured age, gender, and maternal education level. The rationale for the inclusion of specifically these factors lies in their documented correlation with CAPS Scores (Bell et al., 2005). Attention checks that instructed participants to choose a specific answer option were implemented throughout the questionnaires. Participants that failed to select the instructed answer option more than once were excluded.

### 2.5 Procedure

Participants began the experimental session with the described demographics questionnaire. Prior to the instructions, participants were required to position themselves 50cm from the screen and adjust a visually displayed box to match the size of a credit card to ensure that visual stimuli were presented at approximately the same retinotopic size (Li et al., 2020). The main task was then introduced on a component-by-component basis which was inspired by another recent study using a comparable paradigm (Dijkstra et al., 2023). Participants were separately introduced to the two cue dimensions before these were combined and practiced until task difficulty was reached. To ensure the learning of the cue-stimulus-associations, the cues were 100% valid during practice. Additionally, to provide participants with as much practice as possible with the orientation discrimination component, practice trials contained a larger proportion (70%) of grating-present trials than the test session (50%). Participants were informed that these were adjustments to help acquaint them with the task and that these would be reversed in the test phase of the task. They also received feedback and were required to meet certain performance criteria to progress through this training phase. Then, to ensure that the inflated cue validity during the training phase had not introduced a response bias, participants completed one run of trials in which cues were removed and participants again had to reach a performance level of 75%. Finally, participants completed the main task at full difficulty level for 480 trials split into four blocks of 120 trials with 30-second breaks in between. Before finishing the study, participants completed the CAPS and BDI. The testing phase took 40 minutes in total resulting in an overall session length of around 60 minutes including task practice and questionnaires.

### 2.6 Data Analysis

Orientation cues were classified as congruent or incongruent depending on whether the subsequent grating was in line with the cue. Factorized by the presence cues, trials were thus divided into four conditions: presence/absence-cued × orientation-congruent/incongruent. Behavioural analyses compared these four conditions in terms of confidence responses (detection) and orientation response accuracy (discrimination). Results were illustrated using ggplot2 after Allen et al. (2021).

A Bayesian decision model (BDM) was used to further dissect precisely how cues affected perceptual inference. BDMs can be likened to signal detection theory models in the sense that they distinguish between biases and sensitivity-related influences on perception. BDMs have the advantage that they more closely follow the tenets of Bayesian inference and were thus deemed more suitable in this case. The used BDM was adapted from a study conducted by Stuke and colleagues (2021) and modified to fit the present task’s experimental stimuli. The model used participants’ behavioural responses to infer prior and sensitivity parameters. The sensitivity parameter encoded to what extent a participant’s response depended on the displayed stimulus. Based on it, a likelihood value was computed that represented the probability of reporting a certain response option given a trial’s stimulus. The prior parameter on the other hand was a simple free parameter that encoded a shift towards either of the response options regardless of the presented stimulus. Combining the prior with the likelihood, the model produced a posterior probability that the displayed stimulus was correctly chosen, which could be compared to the true behavioural responses on each trial (see Stuke et al., 2021 for exact formulae). This model was then inverted given a large number of trials to estimate the prior and sensitivity parameters by maximizing the summed log-likelihood of predicted responses using Powell’s optimization as implemented in R’s minqa package (Bates et al., 2022; R Core Team, 2022). By comparing the different cueing conditions in terms of the retrieved parameters, the found behavioural effects were further investigated. Specifically, we used the BDM to model detection responses and predicted whether a participant would report a grating as opposed to noise. To this end, confidence responses were binned into low and high relative to each participant’s mean confidence to match the binary input variable of grating or noise. The sensitivity therefore represented the extent to which a participant’s confidence response depended on a grating or noise being shown. The prior parameter reflected biases towards reporting gratings or noise independent of the displayed stimulus.

To assess the role of presence and content cues in hallucination-susceptibility, participant-specific presence and content cue effects were quantified as standardized slope estimates from simple linear regression models predicting confidence and accuracy in individual participants. Linear models were created that predicted participants’ CAPS sum scores from these behavioural effect sizes, Separate models were created to investigate the relation of the two respective cues with CAPS to avoid multicollinearity problems. Each model additionally contained BDI scores as well as the demographic variables age and maternal education level to control for their influences on CAPS scores as rationalized earlier. A square-root transformation was applied to CAPS scores to account for the positive skew in its distribution, in line with past studies (Davies, Teufel & Fletcher, 2017). To allow for comparisons between the types of effects and their relation with CAPS scores, slope estimates from the respective models were standardized. The two samples were well-matched in terms of psychiatric scores and demographics (Table 1).

**Table 1.** Demographics Across the Two Samples. Note: MEL = Maternal education level, CSE = Completed secondary education.

|  | **Age**<br>Mean (SD) | **Gender**<br>(M/F/O %) | **MEL**<br>Mode | **CAPS**<br>Mean (SD) | **BDI**<br>Mean (SD) |
| --- | --- | --- | --- | --- | --- |
| **Original** | 28.2 (5.1) | 51 / 45 / 4 | CSE | 46.5 (49.9) | 13.5 (12.4) |
| **Replication** | 27.1 (6.0) | 64 / 35 / 1 | CSE | 43.2 (49.1) | 13.4 (13.2) |

### 2.7 HOSS simulations

Finally, we considered how our behavioural effects may be accounted for within the framework of the previously described higher-order state space (HOSS) model (Fleming, 2020). HOSS can be specified as a probabilistic graphical model in which *W*is a 1 × *N* vector that encodes the relative probabilities of each of *N* discrete perceptual states (here, left tilted, right tilted, and absent), and *A* is a scalar “awareness state” that encodes the probability of a perceptual state *W*being “present” (left tilt, *w*_1_; right tilt, *w*_2_) or “absent” (*w*_0_). To simulate multivariate sensory data (*X*), *W*in turn determines the value of μ, which is a *M* × *N* matrix defining the location (mean) of a multivariate Gaussian in a feature space of dimensionality *M*. Here we set *M* = 2 to simulate activations of neural populations tuned to the two grating orientations in the task. The means (μ) and covariance (Σ) of *X* were set arbitrarily to roughly match the average accuracies seen in the experiment (μ = [0.5 1.5; 1.5 0.5; 0.5 0.5]; Σ = [0.4 0; 0 0.4]). The priors on the *A* and *W*levels were set to be the same as the empirical probabilities used in the current experiment (*p*(*A*) = 0.2 or 0.8; *p*(*W*) = [0.25 0.75] or [0.75 0.25]).

We drew samples of *X* for different stimuli on “present” trials (mimicking either left or right tilted gratings), under the different combinations of priors at the *A* and *W*levels. 60 samples were drawn for each combination of prior per simulated subject, and for each sample the model was inverted to record posteriors over *W* and *A* as per the following equations using exact inference in Matlab (R2023a):

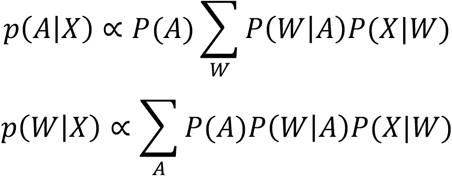

The first equation yields a probability in a stimulus being present vs. absent – this quantity was used to model confidence in having seen a grating in the current experiment. The second equation yields a belief as to which perceptual state was more probable – the maximum of *P*(*w*_1_|*X*) or *P*(*w*_2_|*X*) determines whether the agent responds “left tilt” or “right tilt” to the discrimination decision. Simulations therefore yielded mean confidence and accuracy scores on grating-present trials for 91 simulated individual participants (original sample size post-exclusion), which were analysed in the same manner as the real data.

Initial simulations of this “standard” version of the HOSS model were unable to reproduce the robust presence × content cue interaction effects seen on accuracy in the empirical data. We therefore explored post-hoc alterations to the model architecture to ask what may be needed to capture this salient feature of the data. One attractive model extension (explored in the discussion section of Fleming, 2020) is to allow the sensory covariance Σ to itself depend on the prior over *A* - effectively increasing the range of (content) signals that would be expected under a presence (vs. absence) prior. We implemented this by multiplying Σ by 1.5 when presence was expected (when *p*(*A*) = 0.8).

## 3 Results

### 3.1 Demographics

As previously mentioned, data collection was split into a discovery and replication sample. This split will be further discussed when discussing the hallucination-related analyses, but since no difference between the samples was found in terms of overall behavioural effects, their results will be reported alongside one another when describing the main experimental results. Figures accompanying the text display the original sample unless otherwise specified. Corresponding figures for the replication sample can be found at the end of the results section. No participant in either sample was excluded due to failed attention checks. One participant was excluded in the original sample for failing to respond on a large proportion (29.8%) of trials. Additionally, the described analyses were conducted after excluding 18 and 52 participants respectively based on exclusion criteria aimed at preventing response biases. Ninety-one and 166 participants therefore remained in the original and replication sample respectively. Importantly, these exclusion criteria did not affect the main behavioural results reported here, which indeed remained highly significant in the full sample. Therefore, the exclusion criteria and their practical relevance are restricted to the hallucination-related analyses, in which context they will be discussed in more detail. Behavioural results without excluding any participants can be found in the supplementary materials.

### 3.2 Behavioural results

#### 3.2.1 Presence cues enhance the effect of content cues on discrimination

We first tested whether confidence in having seen a grating tracks the accuracy of discriminating its content. Here we found accuracy to increase linearly with confidence in grating presence, Greenhouse-Geisser-corrected repeated-measures ANOVA: F(1.759, 63.311) = 27.099, p < 0.001, η^2^ = 0.75 (replication: F(1.614, 122.667) = 98.784, p < 0.001, η^2^ = 1.30).

A two (Presence Cue: Present, Absent) by two (Content Cue: Congruent, Incongruent) by three (Contrast: low, medium, high) ANOVA, with repeated-measures on all three factors, was used to analyse discrimination accuracy on grating-present trials. Grating contrast significantly affected accuracy, F(2,180) = 125.605, p < 0.001, η^2^ = 0.58 (replication: F(2,330) = 206.045, p < 0.001, η^2^ = 0.56), so that participants were better at discriminating the content of high-contrast than low contrast gratings, b = 0.141, t(90) = 14.329, p < 0.001 (replication: b = 0.136, t(165) = 17.206, p < 0.001). Importantly, a significant effect of orientation cue congruency was found, F(1,90) = 35.962, p < 0.001, η^2^ = 0.29 (replication: F(1,165) = 35.242, p < 0.001, η^2^ = 0.18), which was grounded in accuracy being higher on orientation-congruent compared to orientation-incongruent trials, b = 0.047, t(90) = 5.997, p < 0.001 (replication: b = 0.05, t(165) = 5.936, p < 0.001), as revealed by post-hoc contrasts. In line with the notion of weighted inference, this effect was found to be stronger on low contrast than on high contrast trials, b = 0.082, t(90) = 4.789, p < 0.001 (replication: b = 0.070, t(165) = 5.161, p < 0.001).

Presence cues did not significantly affect discrimination accuracy, F(1,90) = 1.488, p = 0.226, η^2^ = 0.02 (replication: F(1,165) = 0.707, p = 0.401, η^2^ < 0.01), but they did modulate the effect of orientation cue congruency, as revealed by an interaction between presence and orientation cues, F(1,90) = 11.395, p = 0.001, η^2^ = 0.11 (replication: F(1,165) = 6.559, p = 0.011, η^2^ = 0.04). Further contrast analyses investigating this interaction effect revealed that the effect of orientation cue congruency was moderated so that it was stronger following a presence cue than after an absence cue, b = 0.050, t(90) = 3.376, p = 0.001 (replication: b = 0.030, t(165) = 2.561, p = 0.011). As a result, presence cues only lead to higher accuracy when accompanied by a congruent orientation cue, b= 0.034, t(90) = 3.990, p < 0.001 (replication: b = 0.019, t(165) = 3.388, p < 0.001), but not when accompanied by an incongruent orientation cue, b = -0.016, t(90) = 1.260, p = 0.211 (replication: b = -0.011, t(165) = -1.139, p = 0.256).

A similar pattern of results also appeared for the orientation responses on grating-absent trials. This included an orientation cue effect, F(1,90) = 33.714, p < 0.001, η^2^ = 0.27 (replication: F(1,165) = 42.492, p < 0.001, η^2^ = 0.20), the lack of a presence cue effect, F(1,90) = 0.814, p = 0.369, η^2^ = 0.09 (replication: F(1,165) = 0.315, p = 0.576, η^2^ = 0.02), as well as a significant interaction between the two, F(1,90) = 12.770, p < 0.001, η^2^ = 0.12 (replication: F(1,165) = 17.436, p < 0.001, η^2^ = 0.10). Critically, this also applies to trials on which participants falsely reported seeing a grating with high confidence (Figure 4-B), orientation cue effect: F(1,86) = 41.929, p < 0.001, η^2^ = 0.33 (replication: F(1,157) = 66.850, p < 0.001, η^2^ = 0.30), the lack of a consistent presence cue effect: F(1,86) = 6.190, p = 0.015, η^2^ = 0.07 (replication: F(1,157) = 1.715, p = 0.192, η^2^ = 0.01) and a significant interaction between the two: F(1,86) = 8.312, p = 0.005, η^2^ = 0.09 (replication: F(1,157) = 35.655, p < 0.001, η^2^ = 0.19).

**Figure 4.**
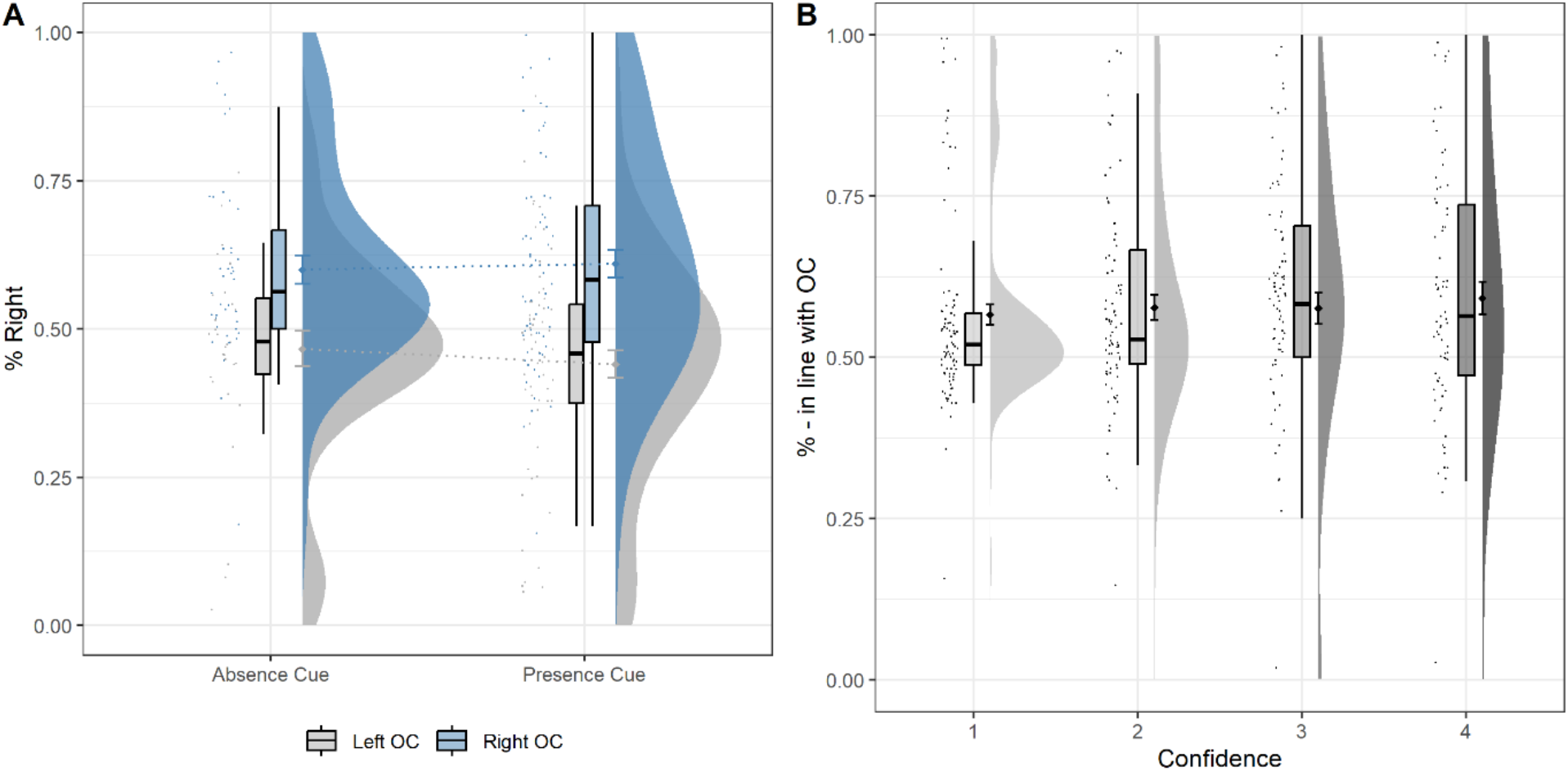
Cue effects on orientation responses on grating-absent trials. (A) Proportion of rightward orientation responses across cueing conditions (OC = orientation cue) (B) Proportion of orientation responses given in line with the trial’s orientation cue (e.g. rightwards response after rightwards cue) across levels of confidence in having seen a grating. Error bars reflect within-subject standard error of the mean.

#### 3.2.1 Content and presence cues independently affect detection

Confidence ratings were similarly analysed on grating-present trials using a two (Presence Cue: Present, Absent) by two (Content Cue: Congruent, Incongruent) by three (Contrast: low, medium, high) ANOVA, with repeated-measures on all three factors (Figure 5-B). Like accuracy, confidence increased with grating contrast, F(2,180) = 275.623, p < 0.001, η^2^ = 0.75 (replication: F(2,330) = 363.448, p < 0.001, η^2^ = 0.69) with participants being more confident in having seen high-contrast gratings than low-contrast gratings, b = 0.656, t(90) = 21.538, p < 0.001 (replication: b = 0.619, t(165) = 23.112, p < 0.001). A significant effect of orientation cue congruency was also once again found, F(1,90) = 33.965, p < 0.001, η^2^ = 0.27 (replication: F(1,165) = 62.842, p < 0.001, η^2^ = 0.25), with congruent cues leading to higher confidence than incongruent ones, b = 0.087, t(90) = 5.825, p < 0.001 (replication: b = 0.265, t(165) = 7.927, p < 0.001). As before, this effect was stronger on low than on high contrast trials, b = 0.102, t(90) = 2.357, p = 0.021 (replication: b = 0.215, t(165) = 10.001, p < 0.001).

**Fig 5.**
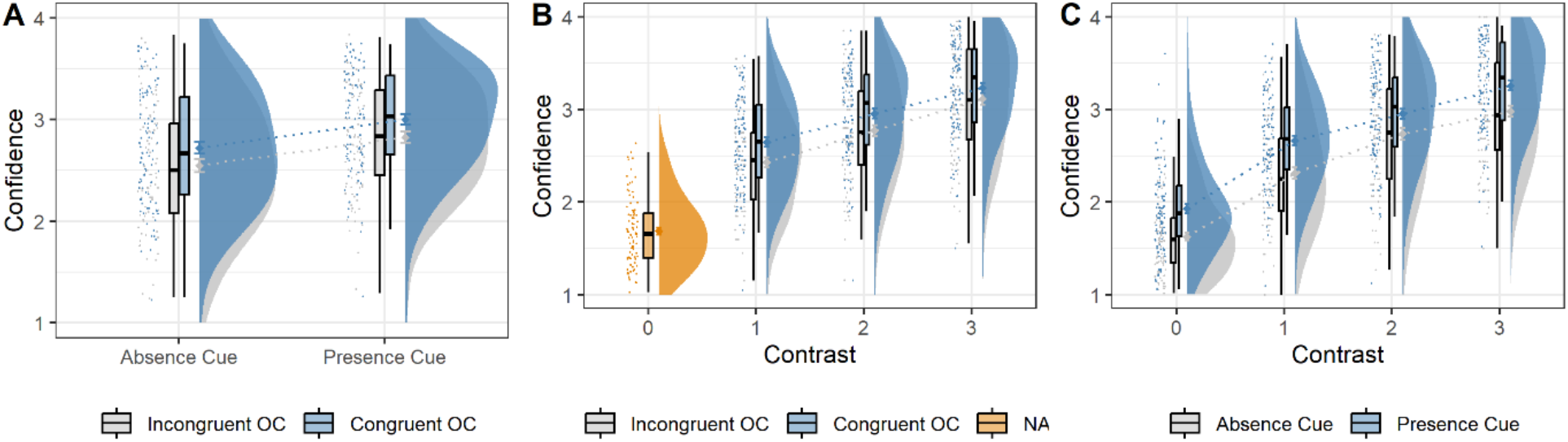
(A) Cue effects on confidence on grating-present trials. (B) Orientation cue congruency (‘NA’ in the case of contrast level 0 since no grating was shown to validate or invalidate the cue) and (C) presence cue effects on confidence across contrast levels (note Contrast = 0 refers to noise trials). Error bars reflect within-subject standard error of the mean.

Additionally, a significant effect was also found for presence cues, F(1,90) = 48.634, p < 0.001, η^2^ = 0.35 (replication: F(1,165) = 56.453, p < 0.001, η^2^ = 0.28), which as expected was grounded in confidence ratings being significantly higher following presence cues than absence cues, b = 0.14, t(90) = 6.964, p < 0.001 (replication: b = 0.426, t(165) = 7.513, p < 0.001).

Unlike the other cue effects, this effect was only significantly stronger on low than on high-contrast trials in the replication sample b = 0.072, t(90) = 1.869, p = 0.065 (replication: b = 0.068, t(165) = 2.333, p = 0.021). The effects of the different types of cues on confidence appeared to be independent as no significant interaction was found between the two, F(1,90) < 0.001, p = 0.977, η^2^ < 0.01. Here, a slight deviation was found in the replication where a small interaction was found, F(1,165) = 4.068, p = 0.045, η^2^ = 0.02, driven by slightly stronger orientation cue effects on confidence on absent cued trials. Interestingly, this subtle interaction was reproduced by the HOSS model (see section 3.3). Analyses of absent trials produced similar results, with presence cues leading to higher confidence ratings than absence cues, F(1,90) = 47.131, p < 0.001, η^2^ = 0.34 (F(1,165) = 56.107, p < 0.001, η^2^ = 0.25).

Overall, content and presence cues were found to have distinct and robust effects on accuracy and confidence ratings. Specifically, content cues were found to promote more accurate responding when the cue was congruent with the stimulus. Crucially, this effect was strengthened or weakened depending on whether a presence or absence cue accompanied the content cue. Aside from this interaction, presence cues were not found to affect accuracy. In contrast, both content and presence cues showed independent effects on confidence, in the absence of robust interactions.

### 3.3 HOSS model simulations reproduce empirical effect profiles

Initial explorations of the standard form of the HOSS model, without any dependency of sensory variance on detection prior, were unable to reproduce the pattern of empirical results – namely the robust interaction of presence and content cues on accuracy. Instead, a model in which presence expectations themselves increased the range of signals that were expected (implementing a dependency between the detection prior and sensory variance) was able to reproduce this qualitative pattern. Here we report these simulations as a proof-of-principle that the HOSS model is flexible enough to accommodate the empirical data patterns, and to inform future computational work (see Methods). However, we note that this is a strictly post-hoc interpretation of the data, and should not be taken as evidence in support of the HOSS model.

The simulated data were adjusted to roughly match the empirical data in terms of both mean accuracy (real: 0.825, simulated: 0.833) and confidence (real: 2.950, simulated, adjusted to task scale: 2.419). Simulated discrimination accuracy scores were analysed using a two (Presence Cue: Present, Absent) by two (Content Cue: Congruent, Incongruent) ANOVA with repeated-measures on both factors (Figure 6-A). As in the empirical data, a significant effect of orientation cue congruency was found on accuracy, which was higher on orientation-congruent compared to orientation-incongruent trials. Presence cues had opposite effects depending on cue congruency, leading to higher accuracy with congruent and lower accuracy with incongruent content cues as reflected in the interaction between the two types of cues (all p < 0.001). Confidence scores were again similarly analysed using a two (Presence Cue: Present, Absent) by two (Content Cue: Congruent, Incongruent) ANOVA, with repeated-measures on both factors (Figure 6-B). Significant main effects were again found for presence and content cues, in line with the real data. The simulated data also reproduced the subtle interaction found in the replication sample of stronger content-congruency effects on absence-cued than on presence-cued trials (all p < 0.001).

**Fig 6.**
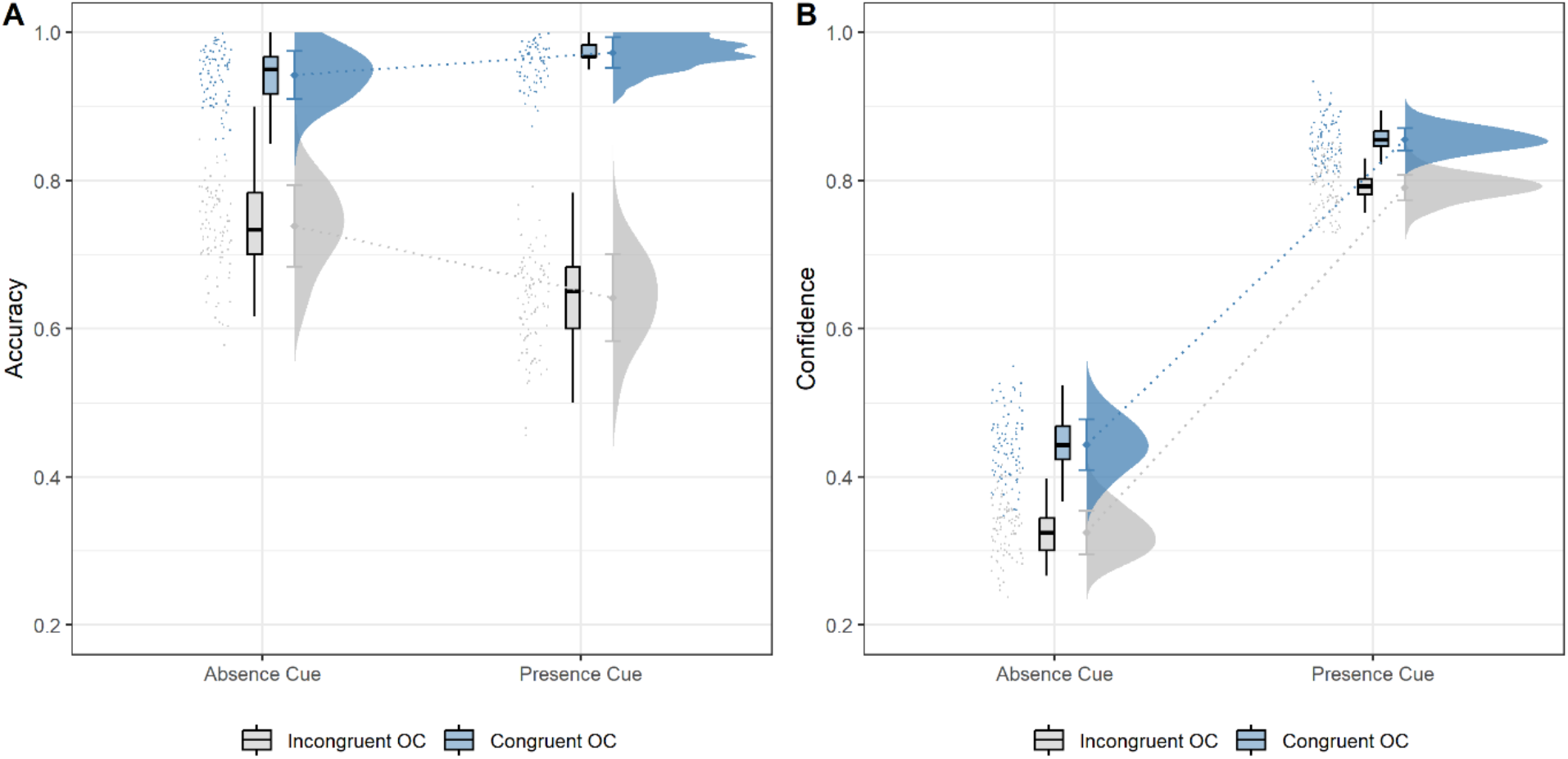
Cue effects on HOSS-model simulated data. (A) Cue effects on accuracy. (B) Cue effects on confidence. Note that for the purposes of modelling, confidence responses were relabelled as 0 (low confidence) and 1 (high confidence). Error bars reflect group-level standard deviation.

In sum, the simulated data were able to reproduce the patterns of empirical effects. This notably includes the contrasting ways in which presence and content cues interact depending on the perceptual decision at hand: presence cues moderate the effect of content cues on discrimination but affect detection in a largely independent manner.

### 3.4 Bayesian decision model results

#### 3.4.1 Presence cues bias detection decisions and affect sensitivity

We further dissected the empirical results using a Bayesian decision model (BDM), to explore how the presence cues affected confidence responses, in particular how presence cues affected the prior probability of reporting high confidence and sensitivity to stimulus presence. A significant difference in retrieved prior parameters was found indicating a bias towards reporting grating presence following presence compared to absence cues, paired t-test: t(90) = 7.432, p < 0.001, d = 0.78 (t(165) = 7.865, p < 0.001, d = 0.61) (Figure 7-A & B, left panel).

**Fig 7.**
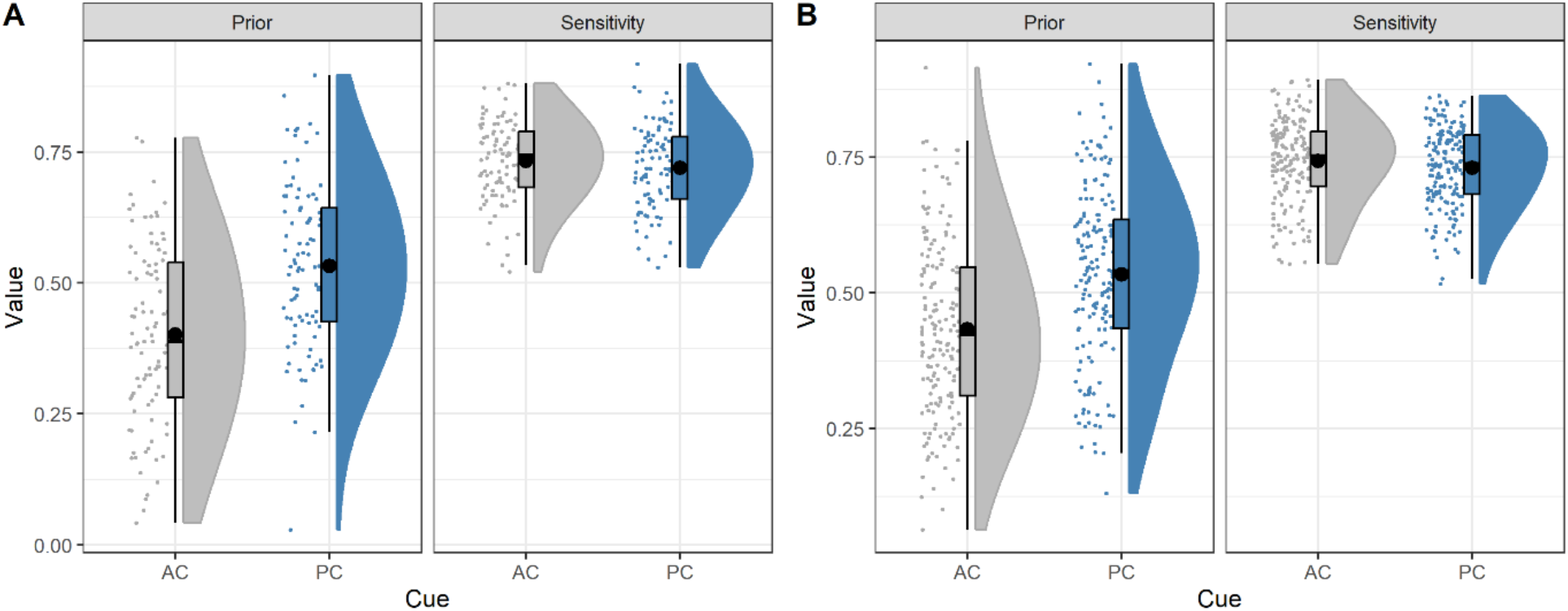
BDM-retrieved parameters for original (A) and replication (B) sample data displayed using the sdamr-package in R (Speekenbrink, 2022).

**Fig 8.**
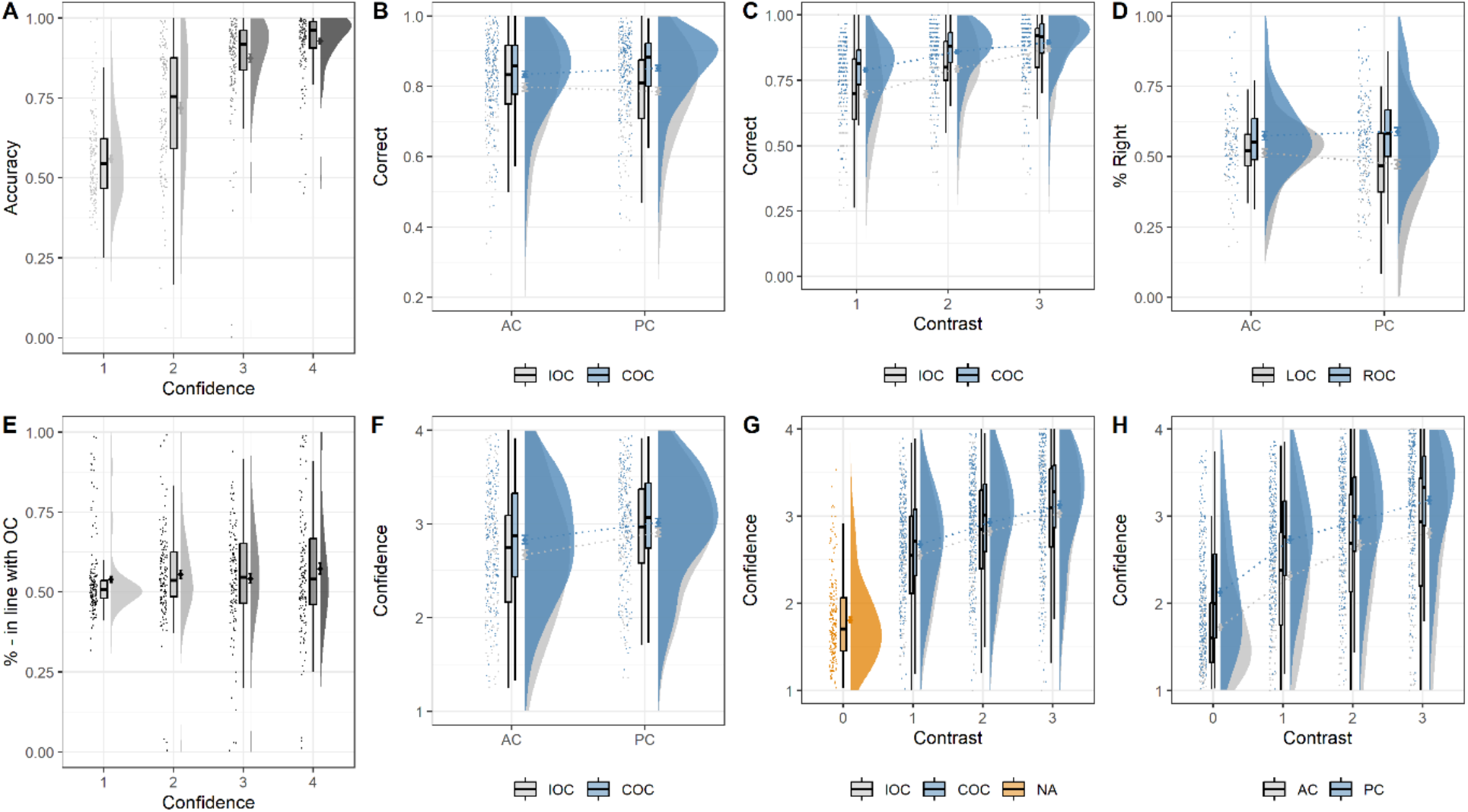
Replication results. Orientation response accuracy on grating-present trials (A) across confidence levels and (B) across cueing conditions (IOC = Incongruent orientation cue, COC = Congruent orientation cue). (C) Effect of orientation cue congruency on accuracy across contrast levels. (D) Proportion of rightward orientation responses on grating-absent trials across cueing conditions (LOC = Left orientation cue, ROC = Right orientation cue, AC = Absence Cue, PC = Presence Cue). (E) Proportion of orientation responses on grating-absent trials given in line with the trial’s orientation cue (e.g. rightwards response after rightwards cue) across levels of confidence in having seen a grating. (F) Cue effects on confidence on grating-present trials. (G) Orientation cue congruency and (H) presence cue effects on confidence across contrast levels.

Interestingly, a significant effect of presence cues was also found with regards to the retrieved sensitivity parameters, however in the opposite direction, t(90) = 2.193, p = 0.03, d = 0.23 (t(165) = -2.955, p = 0.004, d = 0.23), meaning that higher sensitivity parameters were found for the absence-cue compared to the presence-cue condition (Figure 7-A & B, right panel). In other words, participants’ confidence in stimulus presence more accurately tracked the actual presence of a grating following an absence cue.

### 3.5 Hallucination-susceptibility results

In the original sample, the content cue effect was not found to predict CAPS scores, β = 0.008, t(85) = 0.080, p = 0.936 while a marginal effect was found for the presence cue effect, β = 0.166, t(85) = 1.784, p = 0.078, η^2^ = 0.04. However, the trend-level relationship between the presence cue effect and CAPS was not present in the replication sample, β = 0.024, t(161) = 0.366, p = 0.715, and neither was the content cue effect, β = 0.073, t(161) = 1.126, p = 0.262. The results were obtained after applying the following exclusion criteria. Participants were excluded if they (1) performed significantly worse than chance on orientation-incongruent trials or (2) showed no effect of grating presence on confidence responses. The rationale behind both criteria was to exclude participants who did not respond based on the actual stimuli on the screen. Interestingly, relaxing these exclusion criteria resulted in a slightly stronger association between the presence cue effect and CAPS across the original and replication samples: β = 0.173, t(104) = 2.066, p = 0.041 (β = 0.095, t(213) = 1.615, p = 0.108). This suggests that any relationship between the presence cue effect and hallucination-susceptibility was driven by a subset of participants whose responses were particularly driven by the cues. Supporting this notion, an ad-hoc leave-one-out analysis, which can be found in the supplementary materials, showed the effect of interest to be strongly reliant on individual participants.

## 4 Discussion

Across two independent samples, this study introduced and validated a novel design for studying separate effects of presence and content priors on perceptual decision-making. The results provide robust empirical support for the distinct effect profiles of these different kinds of predictions. Specifically, our findings are consistent with more global presence predictions gating the effect of content predictions on lower-level discrimination responses. Meanwhile, presence and content predictions independently influenced higher-level detection. By informing a theoretical distinction between predictions of presence and content with empirical evidence, these findings illustrate the potential value or even necessity of introducing more nuance into the construct of prediction as well as its investigation.

Beyond their distinct effects, our study also demonstrated the interaction between presence and content priors. Specifically, the biasing influence of content priors on discrimination judgements was boosted in the case of a presence prior and dampened in the case of an absence prior. This follows intuition in the sense that the utility of any content-specific prediction is scaled by the likelihood of a stimulus appearing in the first place: the more likely the answer to the question ‘will something appear?’ is no, the more preconceptions about ‘what will it be?’ become redundant. Interestingly, a recent neuroimaging study also found an effect of content cue congruency on prefrontal activity to be contingent on the expectation of general stimulus presence (Dijkstra et al., 2023). This suggests that presence expectations may act as a regulatory volume knob for the influence of content predictions lower down the hierarchy.

From a computational point of view, while standard signal detection theory models of detection and discrimination frame the two as evidence accumulation within the same sensorial landscape rendering them both first-order processes (King & Dehaene, 2014), the flat structure of such a model does not intuitively provide room for how hierarchical effects of one type of prediction affect the other. One computational model has put forward that these different kinds of predictions have a hierarchical relationship to one another (Fleming, 2020), which we here adopted and modified to demonstrate that the behavioural effects in our paper could at least in principle be accounted for by such a model. Note however, that this is not evidence for the brain deploying such a hierarchical predictions, as the model was adjusted to fit our data. Separate future empirical studies will need to provide evidence for the existence of such a computational hierarchy.

The current study also provides insight into more precisely how presence priors affect perceptual inference. Specifically, they led to biases in detection judgements, as expected based on past literature (Wyart, Nobre & Summerfield, 2012). However, the impact of presence priors on detection was also accompanied by a loss in overall sensitivity to sensory evidence. This suggests that participants were better at distinguishing between gratings and noise following an absence cue than a presence cue. One potential explanation for this unexpected finding is that after an absence cue, participants start with an expectation of noise, and need to accumulate sensory evidence in favour of grating presence to make a correct decision on grating-present trials. In contrast, following a presence cue, participants already expect to see a grating and need to gather evidence for its absence to make a correct decision on grating-absent trials. Past research has shown that an asymmetry exists between gathering evidence for absence vs presence, such that it is easier to gather evidence for presence (Mazor et al., 2019). This is thought to be due to absence decisions requiring negative evidence – something not being there – whereas sensory systems are typically tuned to propagate positive evidence for something being there (Meuwese et al., 2014). The result is that participants may be naturally better at accumulating evidence on incongruent absence-cued trials than incongruent presence-cued trials, leading to the difference in sensitivity parameters. Hierarchical models additionally suggest that prediction errors regarding stimulus presence are abstract, and lack content-specific information (Dijkstra et al., 2023; Fleming, 2020). This may explain why, despite increased sensitivity to stimulus presence, accuracy on orientation responses was not higher following absence than presence cues. It should be noted that while supported by the current findings, this specific explanation is at this point largely speculative and requires further empirical investigation.

Having successfully established the distinct ways in which presence and content priors affect normative perceptual decision-making, this study sought to further elucidate their roles in hallucination-like perception, which has previously been attributed to an overly strong reliance on predictions (Powers et al., 2016; Sterzer. et al., 2018; Corlett et al., 2019). Indeed, in the present study we found that on grating absent trials, high confidence false percepts were reported in line with the cued orientation. Moreover, expecting the presence of a stimulus increased the chance of detecting one on grating-present trials. In previous work where participants were unaware of the purpose of the cues, we found that content cues did not necessarily induce false percepts. However, in these studies false percepts still arose through stimulus-like signals reflecting the falsely perceived orientation (Haarsma et al., 2023; Haarsma et al., 2024). One possibility is that explicit content cues induce sensory-like activity (Kok et al., 2017; Aitken et al., 2020) resulting in false percepts.

We next tested whether the strength of content and presence predictions in driving false percepts related to hallucination-like perception. In both samples, there was no evidence for a relationship between the strength of content predictions and hallucination-proneness. This is in line with previous studies where content cues were not associated with hallucination-like perception (Haarsma et al., 2023). In contrast, in the discovery sample we did find that presence predictions have an enhanced influence on confidence in stimulus presence on absent trials, suggesting that presence priors are indeed overly strong. However, this result was not replicated in a second, larger sample, and largely dependent on exclusion criteria, specifically the inclusion of participants that were heavily influenced by the cues. This can hinder a straightforward interpretation of the results, as these participants may have disregarded the stimuli altogether. This issue is largely unaddressed in previous work, but has particular relevance for studies employing explicit cue instructions. The lack of an overly strong prediction effect in people prone to hallucinations in the replication sample seemingly contradicts the past literature. Although some past studies have found this type of effect in similarly sized or smaller, normative samples (Haarsma et al., 2023; Stuke et al., 2021), larger sample sizes may be required to observe these effects consistently. Furthermore, it is important to note that the average CAPS scores in this study were considerably lower (averaging around 45, in line with the original study by Bell et al, 2005) compared to some previous studies, which reported average CAPS scores of 107 (Stuke et al., 2021), 72 (Haarsma et al., 2023) and 58 (Schmack et al., 2021). Our samples’ limited range of hallucination-like experience might have reduced the power of this study. Future studies could enrich their samples by targeting clinical groups.

In summary, our study aimed to empirically establish a distinction between presence and content priors by teasing apart their influences on perceptual decision-making. Beyond demonstrating that these priors differentially affect detection and discrimination decisions, we also showed that presence priors scale the influence of content priors on discrimination judgements. This is consistent with the proposal of a hierarchical relationship between these two priors, which an updated HOSS model was able to capture. Together, the current findings shed light on the computational underpinnings of the influence of expectations on perceptual inference.

