## Supplementary Material for "Expectations about presence enhance the influence of content-specific expectations on low-level orientation judgements"

1. Full results without exclusion

a) Original sample

Discrimination Accuracy :

Orientation cue congruency: $F(1,109) = 43.547$, $p < 0.001$
Presence cue: $F(1,109) = 2.358$, $p = 0.128$
Orientation cue congruency X Presence cue: $F(1,109) = 10.210$, $p = 0.002$
Contrast: $F(2,218) = 121.209$, $p < 0.001$
Contrast X Orientation cue congruency: $F(2,218) = 9.541$, $p < 0.001$

Confidence :

Orientation cue congruency: $F(1,109) = 40.442$, $p < 0.001$
Presence cue: $F(1,109) = 38.898$, $p < 0.001$
Orientation cue congruency X Presence cue: $F(1,109) = 0.395$, $p = 0.531$
Contrast: $F(2,218) = 201.108$, $p < 0.001$
Contrast X Orientation cue congruency : $F(2,218) = 4.768$, $p = 0.009$
Contrast X Presence cue : $F(2,218) = 13.474$, $p = 0.001$

b) Replication sample

Discrimination Accuracy :

Orientation cue congruency: $F(1,217) = 66.587$, $p < 0.001$
Presence cue: $F(1,217) = 0.661$, $p = 0.417$
Orientation cue congruency X Presence cue: $F(1,217) = 14.528$, $p < 0.001$
Contrast: $F(2,434) = 143.302$, $p < 0.001$
Contrast X Orientation cue congruency: $F(2,434) = 11.700$, $p < 0.001$

Confidence :

Orientation cue congruency: $F(1,217) = 69.072$, $p < 0.001$
Presence cue: $F(1,217) = 79.604$, $p < 0.001$
Orientation cue congruency X Presence cue: $F(1,217) = 2.793$, $p = 0.096$
Contrast: $F(2,434) = 237.19$, $p < 0.001$
Contrast X Orientation cue congruency : $F(2,434) = 4.221$, $p = 0.015$
Contrast X Presence cue : $F(2,434) = 15.517$, $p = 0.001$

2. Leave-one-out analysis

To illustrate the dependency of found effects on individual subjects, a leave-one-out analysis loop was devised that repeated the aforementioned analysis relating cue effects to hallucination susceptibility while excluding one participant from the analysis on every run. Figure S1-A shows the slope estimates for the higher-level cue effect's relation to CAPS scores on every run of this analysis. What becomes evident is that while a general centring around the group-level effect is apparent, the exclusion of single participants has rather drastic effects on the found effect. In fact, the exclusion of one particularly influential participant (marked) was found to lead to a slope estimate three times as large as if no participant was excluded.

Applying the same analysis to the original sample, where the trend-level effect was found that the replication study was based on, yielded a similar pattern of results (Figure S1-B). The exclusion of a single participant was found to eradicate the found effect. For comparison, the relationship of the covariate MEL with CAPS which is comparable in its effect size was not found to be as dependent on particular participants (Figure S1-C). This variability in effects based on the exclusion of participants was not specific to the used parametric models either as non-parametric measures such as Spearman correlations showed an analogous pattern. Overall, while not necessarily surprising, this dependence of results on individual participants may serve as a call for caution in these types of analyses relating behavioural measures to psychiatric questionnaires.

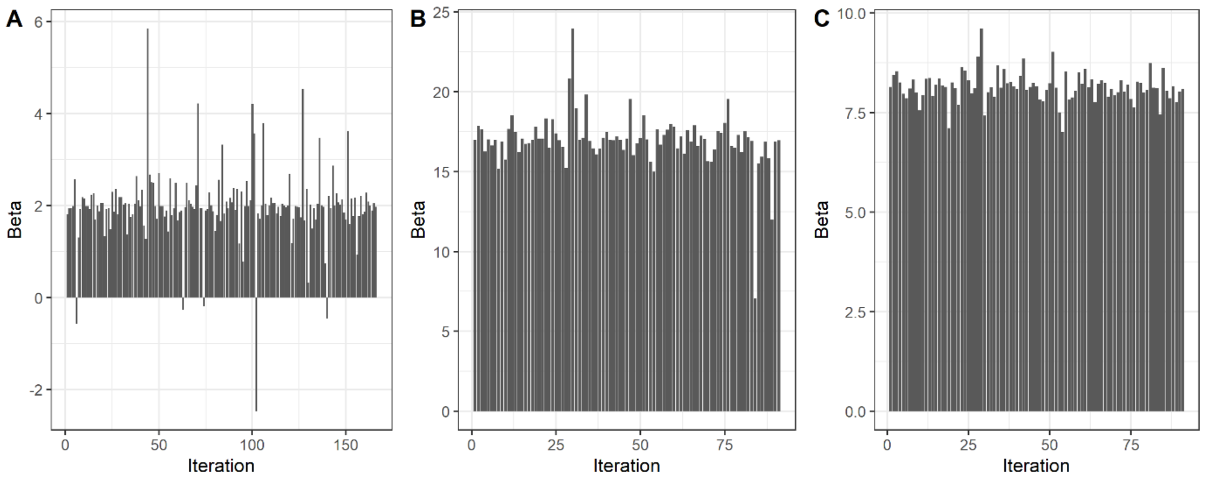


Fig 5. LOO-Analysis results for the effect of interest in the (A) replication and (B) original sample. (C) For comparison, the same LOO-analysis applied to the effect of MEL on CAPS in the original sample, matched in effect size with (B), $\eta^2 = 0.06$.
